# Intraspecies variability in plant and soil chemical properties in a common garden plantation of the energy crop *Populus*

**DOI:** 10.1101/2023.06.16.545338

**Authors:** Matthew E Craig, Anne E Harman-Ware, Kevin R Cope, Udaya C Kalluri

## Abstract

Optimizing crops for synergistic soil carbon (C) sequestration represents a frontier approach toward CO_2_ removal in food and bioenergy production systems. While the central roles of plants in biological C capture and storage belowground in soils is well known, we lack an understanding of how intraspecies variation in bioenergy plants affects soil biogeochemistry. This knowledge gap is exacerbated by spatial heterogeneity in soil and plant systems, and by the difficulty of characterizing belowground plant traits. Here, we sought to obtain first insights on the spatial variation of C and nutrients in soil and plant tissues from a common garden field site of diverse, natural variant, *Populus trichocarpa* genotypes—grown and characterized previously for aboveground biomass-to-biofuels research. Such field sites represent a potential resource for evaluating genotype-specific effects on soil C, but this usage may be complicated due to dense plantings of intermixed genotypes. Thus, we sampled soils at the scale of individual trees to determine whether it is feasible to detect soil property variation with different plant genotypes in this system. We additionally sampled stem and root tissues to evaluate the potential for inferring important belowground traits based on aboveground-belowground correlations. We found that substantial variation in soil properties could be explained at the scale of individual trees, suggesting that genetically diverse plantations can be used to assess plant-soil correlations. Though we did not observe genotype-specific patterns in soil C, other properties such as soil acid-base chemistry (soil pH and base cations) and bulk density showed genotype-specific correlations. Stem and root nutrient levels were generally not correlated, suggesting that belowground traits should be measured directly. In conclusion, our pilot study suggests that long-term common gardens of genome-wide association study populations represent useful resources for understanding plant genotypic relationships with soil properties in *Populus* field study test plots. These resources could be used to develop verified plant species, geographic region-specific standardized sampling methods, and baseline data. Such context-specific, empirically verified data and models will be necessary for informing applied research strategies in selecting high aboveground productivity genotypes for enhanced soil C storage in managed, commercial scale, woody bioenergy crop plantation systems.

## Introduction

Enhanced soil carbon (C) storage—i.e. removing CO_2_ from the atmosphere in favor of a longer-lived terrestrial pool—represents one possible nature-based climate solution (Lal, 2004; Minasny et al., 2017). Agricultural systems—including bioenergy systems—have the greatest potential for enhanced soil C because these systems have been the most depleted due to human land use (Davidson & Ackerman, 1993; Sanderman et al., 2017). Frontier technologies offer the potential to enhance soil C sequestration beyond what is currently achievable with best management practices (Paustian et al., 2019), especially if combined with other strategies like BECCS (bioenergy with carbon capture and storage) (Field et al., 2020). One promising frontier approach is to optimize crops for synergistic soil C sequestration (Paustian et al., 2016). This involves identifying candidate genotypes that associate with soil C promoting phenotypes (Yang et al., 2021) or, better, genotypes that associate with positive soil C outcomes (e.g. net accumulation). Detecting genotype-soil C associations is challenging because there are few resources suitable to answer this question and because soil biogeochemistry changes slowly and via many mechanisms in response to plant trait variation.

Long-term (e.g. >10 years old) plant genetic diversity panels, planted in common gardens for genome wide association studies (e.g. Muchero et al., 2018), offer an opportunity for unprecedented studies of associations between plant genotypes and soil C dynamics. Several such plantations exist for the bioenergy candidate, *Populus trichocarpa* (Evans et al., 2014). However, in these systems, genotypes are intermixed and individual trees are planted in 1-2 m apart (Chhetri et al., 2020). Studying effects of individual genotypes on soil properties, therefore, requires a focal tree approach, instead of the plot-level measurements that are typical in ecosystem-level soil biogeochemistry studies. This presents a potential methodological barrier because soil C and other biogeochemical properties are spatially heterogeneous at fine spatial scales (Garten Jr. & Wullschleger, 1999), which can also drive fine-scale variability in plant traits (Harman-Ware et al., 2021). Thus, prior to any broad-scale sampling efforts to leverage and inform plant-soil interactions in large genetic diversity plantation systems, there is a need for empirical evaluation of sampling strategies and to address whether plant-soil couplings are detectable, variable, and replicable at the scale of individual plants or genotypes.

The difficulty of measuring belowground plant traits presents an additional challenge for studying genotype associations with soil C and other properties. Belowground plant inputs, including root tissues and exudates, are now recognized as critical drivers of soil C formation (Keller et al., 2021; Rasse et al., 2005; Sokol & Bradford, 2019; Villarino et al., 2021), and soil microbial traits mediate both soil C formation and loss (Schimel & Schaeffer, 2012). Thus, assessing belowground plant and microbial traits is critical to understanding soil C dynamics, especially in agricultural and managed forest systems where aboveground biomass is harvested. Focal tree root sampling is time consuming in these plantation systems because it requires tracing excavated roots to the genotype of interest. The challenging nature of sampling and measurement has therefore resulted in asymmetric knowledge gaps in belowground dynamics relative to heavily studied above-ground dynamics in bioenergy plantations. Inferring belowground traits from aboveground traits is one potential way around this issue. However, aboveground-belowground trait associations have mixed support among species (Sun et al., 2018), and not much is known about these associations among different field-cultivated genotypes for *P. trichocarpa*. In summary, *Populus* Genome-wide association study (GWAS) plantations represent a promising system for identifying important genotype-specific associations with belowground properties but given the challenges of working in such a system, an initial evaluation is critical to design informed sampling campaigns at scale.

In this study, we worked in a genetically diverse, long-term *P. trichocarpa* population—with a wide range of phenotypic variation reported in aboveground traits—growing in a common garden field setting in the Pacific Northwest, USA (Happs et al., 2021). Belowground soil, plant, and microbial traits have not been studied in this population. While the dense plantings of intermixed genotypes can yield plant--soil linkages at individual trees, replication of genotypes in three replicate blocks provides the opportunity for capturing genotypic effects. Our goals in this first study of a woody perennial bioenergy plant and associated soil on a ten-year old intraspecific GWAS field site were to evaluate 1) the potential of focal tree soil sampling for detecting associations of poplar genotypes with soil biogeochemistry (e.g. C, nitrogen (N), phosphorus (P), and a range of soil cations) and microbial community composition, and 2) the strength of correlations between above- and belowground traits (stem vs root and plant vs soil). To do this, we measured soil, root, and stem chemical traits for several focal trees spanning four different *P. trichocarpa* genotypes representative of population extremes in cell wall chemistry phenotypes low (BESC 24) or high (BESC 375, BESC 371, BESC 394) lignin content based on population-wide cell wall chemistry data previously reported for this population in several prior reports (Studer et al., 2011; Muchero et al., 2015; Happs et al., 2021; Harman-Ware et al., 2021 and 2022). We hypothesized that 1) different *P. trichocarpa* genotypes determine variation in soil properties, and that this variation is observable at the scale of individual trees, 2) plant lignin phenotype influences amount of C in soil, and 3) belowground traits correlate positively with aboveground traits across genotypes. We also explored the influence of sampling location on measured soil properties, potential effects of plant tissue chemistry on soil C, relationships among nutrient elements beyond C in plant and soil pools in bulk soils sampled at conventional organic matter rich, surface soil horizon depth of 0-10 cm in the pilot study.

## Materials and Methods

### Field site description

The *Populus trichocarpa* GWAS Common Garden site field site in Oregon is located in Clastkanie; spatial coordinates: Latitude: 46.1196, Longitude: -123.2629. The site is a flat land with a relatively uniform soil type; Entisols (Veach et al. 2019) formed by floodplain activity adjacent to the Columbia River. Continuous land use in past decades has been for *Populus* plantations. Under the present study, we collected surface soil (top 10 cm) from under the planted and non-planted areas at the site and analysis of basic soil characteristic revealed an average of pH of 5.33, % moisture of 31.3, C:N ratio of 12.24, and soil texture as Silty Clay with 9.97 % sand, 47.5 % silt and 42.5 % clay. The common garden site has a thousand genotypes of *P. trichocarpa*, densely planted (<2 m apart) and replicated in three blocks with randomization and surrounded by two edge rows on all four edges of the field site, and stably growing for over 11 years.

### Field sample collection

Soil samples (0-5 cm [top] and 5-10 cm [bottom] depth) were collected at a radius of one foot from the focal tree trunk, noting cardinal direction, digging using graduated trowels to reference depth and collecting and storing in labeled Ziploc bags. Two replicate trees were sampled for each of the four genotypes. For bulk density analysis, soil samples were collected using AMS Soil Core Sampler Kit with Hammer Attachment (part # 77455), 2 in × 2 in Soil Core Kit and stored in 2 in × 2 in core liners Precut (plastic). Samples were shipped on ice and stored at 4 °C until analysis. Stem cores (1 cm in diameter) from tree breast height were collected using increment borer and stored and shipped in zip-lock bags at −20 °C. Fine roots (<2 mm diameter) were accessed by shallow digging, tracing of roots to tree stump and cut using ethanol wiped pruners and stored and shipped in ziplock bags on dry ice. For elemental analysis, subsamples of roots and stems were separately aliquoted into labeled brown paper bags and dried in hot air oven 70 °C for three days and shipped to soil service lab in University of Georgia.

### Laboratory analyses

Basic soil analysis and elemental analysis of plant and soil samples was conducted at the soil testing service of UGA Extension, Agricultural and Environmental Services Laboratories using standard methods. Soil pH, Lime requirement, Phosphorus, Potassium, Calcium, Magnesium, Zinc, Manganese were performed using published standard methods (Mehlich 1953; Kissel et al. 2012). Soil texture analysis was performed using Hydrometer (Gee and Bauder, 1996). Total Carbon (C) and total Nitrogen (N) was determined using published Soil Science Society of America methods (Bremner, 1996; Nelson and Sommers, 1996)

### Data analysis

To examine the extent to which soil properties vary at the scale of individual trees, we fit linear models with focal tree and sample depth as predictors (R version). Assumptions of normality and homogeneity of variance were checked by inspecting residual plots and with a Shapiro-Wilk test. Response variables were natural log-transformed where necessary to improve adherence to assumptions (specifically for soil C and N content). To determine the proportion of variance explained at the level of individual trees, partial R^2^ were determined using the ‘asbio’ package (Kutner et al., 2005). To provide an initial look at whether soil variables are clustered with respect to genotype, we additionally calculated partial R^2^ for models fit with genotype and sample depth as predictors. However, these analyses are considered exploratory due to inadequate replication, so we do not report other test statistics for genotype scale analyses.

Relationships, principal components and variance amongst stem, root and soil traits (elemental, chemical and physical properties) were measured by determination of Pearson Correlation Coefficients (PCC) and Principal Component Analysis (PCA). PCC values were calculated on raw data values and PCA (100 iterations of NIPALS algorithm, 20 random cross validation) was performed on mean-centered compositional data using the Unscrambler X V.10.5 (Camo Analytics, AspenTech).

## Results and Discussion

We evaluated whether genetically diverse, long-term (ten-year) stands of a natural *Populus trichocarpa* collection, present a practicable resource for quantifying genotype-specific plant effects on soil chemical properties, including soil C (Fig 1A). If so, studying such systems could facilitate the optimization of commercial poplar systems for soil C storage. Our study system is well suited to address this question because soil biogeochemical properties are spatially heterogeneous and change slowly. Our analysis represents a first attempt to evaluate aboveground-belowground correlations among *P. trichocarpa* genotypes and soil properties in a genetic diversity plantation. Given that different genotypes are intermixed within replicated blocks, our analysis informs the extent of feasibility of studying aboveground-belowground correlations in this system. Across the entire field site soil, correlation between soil C and Soil N was strong (r=0.89; Fig. 1B). Correlation analysis of all measured soil parameters (pH, bulk density and elements) provided insights on additional strongly correlated soil properties (Fig. 1C). PCA analyses were also conducted to explore whether there is a dependence of overall soil chemistry on either the radial direction (East, North and West) or depth (top [0-5 cm] vs bottom [5-10 cm]) of sampling, and our results showed no significant effect overall (Suppl Fig. 1). However, with specific soil variables such as pH, Ca, K, Mg, and P (Fig. 2A), linear models did show an association with sample depth, such as where pH and nutrient concentrations were found to higher in the top surface layer. Next, PCA analysis undertaken using one high (BESC 375) and one low lignin (BESC 24) genotype showed genotype-specific clustering of soil properties (Suppl Fig. 2), which was not observed with inclusion data of all four genotypes.

**Fig. 1.**
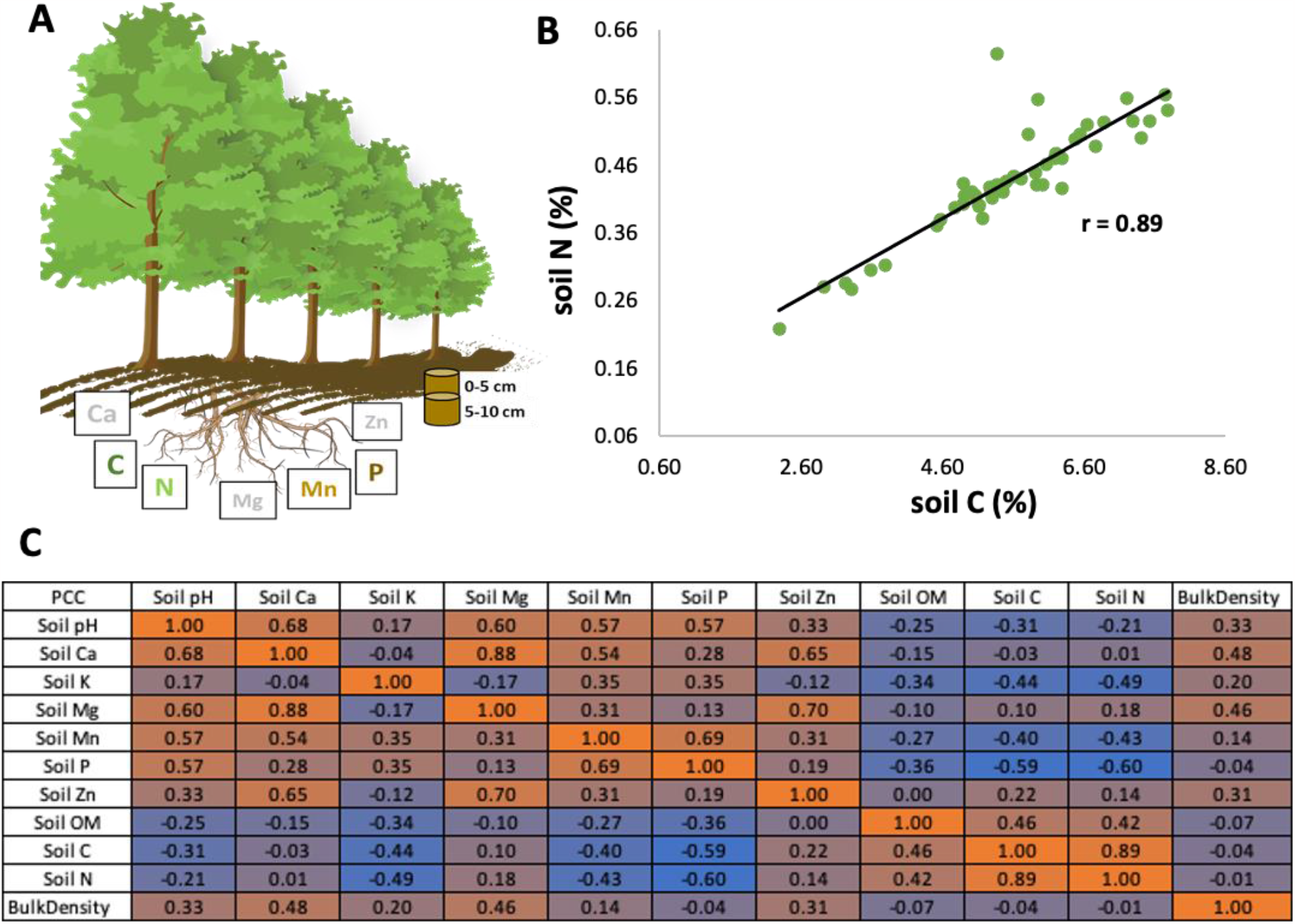
Correlation analysis undertaken at Clatskanie *Populus* common garden site. (A) Graphical representation of plant-soil elemental distribution analysis undertaken in top 10 cm surface soil profile (bulk soil collected from 0-5 cm and 5-10 cm depth pools). (B) Bivariate between C and N using all soil samples (R^2^ = 0.79). (C) Correlation matrix for all soil properties measured.

**Fig. 2.**
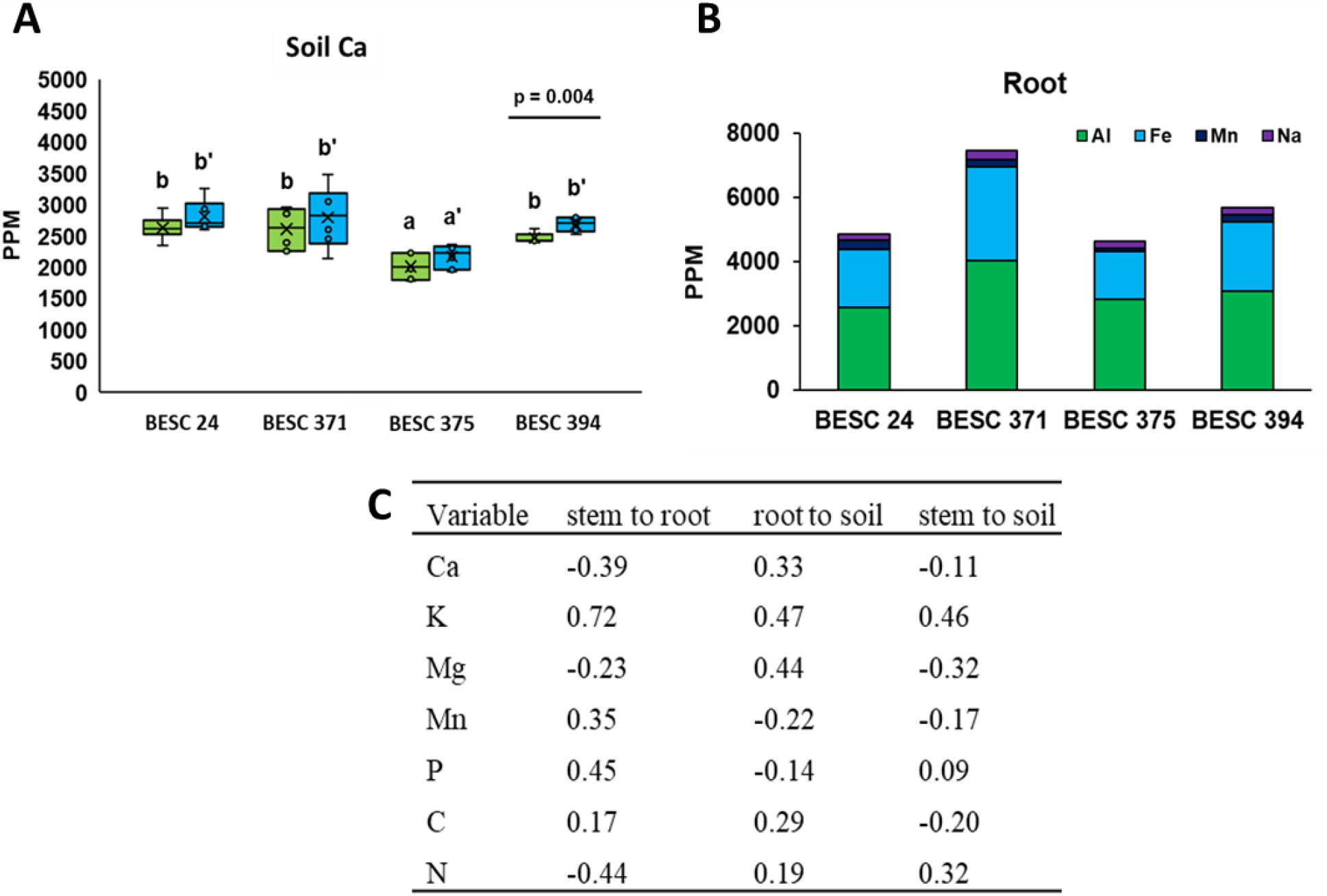
Representative observations of differential correlation of elemental distribution with genotype, within tissue type and across above and belowground samples. (A) Ca level in soil collected from BESC 375 from across the replicate blocks was observed to be significantly different relative to other genotypes based on ANOVA (p < 0.05). Horizontal bars indicate significant differences between top (green) and bottom (blue) soil samples. (B) Within the same plant tissue (root), elemental distribution (shown here Al, Fe, Mn and Na) varied across genotypes. (C) Correlation between above-belowground distribution of elements is low (shown here Pearson’s correlation coefficients between element distributions in stem to root, root to soil stem to soil). A comprehensive correlation matrix generated using all parameters measured for stem, root and soil samples obtained from the four genotypes is provided in Supplemental File.

With more targeted correlation analyses, we observed differential elemental distribution with genotype, within tissue type and across above and belowground samples (Fig. 2). Ca level in soil collected from BESC 375 from across the replicate blocks was observed to be significantly different relative to other genotypes (Fig. 2A). Within the same plant tissue (root), elemental distribution (shown here Al, Fe, Mn and Na) varied across genotypes (Fig. 2B). Correlation between above-belowground distribution of elements, based on Pearson’s correlation coefficients between element distributions in stem to root, root to soil stem to soil samples, were low (Fig. 2C). A comprehensive correlation matrix generated using all parameters measured for stem, root and soil samples obtained from the four genotypes is provided as Suppl. Table 1.

Plants were also associated with differences in soil physical properties. *Populus* presence, regardless of individual, genotype, or sample depth, was associated with significantly lower soil bulk density compared with a control microsite i.e. a location containing no *Populus* individual within a 5 m radius and with no plant roots evident in the soil. In general, nutrient concentrations were not correlated across tissues (stem-root) or between tissues and soil (stem-soil, root-soil; Fig 2C). One possible exception to this is stem and root K concentrations, which were positively correlated (PCC = 0.72). Correlations among different elements within a pool were also detected. For example, soil bivalent cations (Ca^2+^ and Mg^2+^) were positively correlated and were positively correlated with soil pH (Suppl Table 1). Correlations were also observed across pools and elements. For example, soil N was negatively related to root K and stem K.

We found limited evidence to suggest coordination of stem and root traits among plant genotypes (Suppl. Table 1). However, data from the pilot study support that soil properties can cluster at the genotype or individual tree level (Suppl Fig. 3). Focal tree was a significant predictor of all analyzed soil properties (*p* < 0.001), and explained more than half of the variation 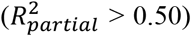 in all properties except for soil C:N (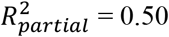; Suppl Table 1). For models using data aggregated at the scale of genotype, variables related to soil acid-base chemistry clustered together for different genotypes. Specifically, genotype predicted more than half of variation in pH and Ca and predicted more than a quarter of variation in Mg and Mn (Suppl Table 1). Visual inspection of plots reveals that one genotype, BESC 375, is a strong driver of these patterns (Suppl Fig. 3). BESC-375 was associated with more acidic (low pH), calcium-depleted soil for both measured individuals. In general, the genotype representing the low end of lignin distribution in the population, BESC 24, was not observed to have specific effects on soil chemical properties. We did note a tendency for BESC-24 to associate with higher bulk densities in the deeper measured soil layer.

In general, our results support our first hypothesis that *P. trichocarpa* genotypes differentially associate with soil biogeochemical properties, and that this is observable at the scale of individual trees. The pronounced variability in soil biogeochemical properties across our common garden indicates that planted trees are likely driving differences in soil C and nutrient cycling, though we note that there could be other sources of soil heterogeneity (including microbial associations), even in a common garden. We found that most of the variation in soil biogeochemical properties in this system occurred among individual focal trees, exceeding micro-scale variability within each focal tree location. Typical common garden studies of tree effects on soil biogeochemistry employ a plot-based approach and are designed to detect changes over a larger area seeded with a common species (e.g. Vesterdal et al., 2008). Fine-scale heterogeneity could make it difficult to detect species effects at a smaller spatial grain. For example, working in natural forests, (Fraterrigo et al., 2005) measured high variability in soil chemical properties within 20 × 20 m plots, but found that previous land use generally had a homogenizing effect at this scale. It is likely that previous land use has also led to reduced fine-scale soil heterogeneity in our study site. Regardless, our finding that soil biogeochemical properties clustered at the scale of our focal tree sampling indicates that our site, and similar plantations, could be used to study genotype-specific effects on soil biogeochemistry.

We also found preliminary evidence that *Populus* genotypes differentially affect soil biogeochemical properties. Specifically, soil pH and soil base cations (except for potassium) were substantially lower for one genotype, BESC 375 (across all replicates distributed across the plantation). A number of traits could lead to differences in acid-base chemistry under different strains of poplar including association with ecto-versus arbuscular mycorrhizal fungi, N or Ca uptake mechanisms, or organic acid production, to name a few (Ehrenfeld et al., 2001; Finzi et al., 1998; Lin et al., 2022). Though we did not find evidence for genotype effects on soil C in this limited sampling effort, we note that these effects on soil acid-base chemistry could indicate or lead to effects on soil C cycling. For example, soil Ca availability could enhance mineral-associated C formation via cation bridging mechanisms (von Lützow et al., 2007), an effect which likely depends on soil pH (Rasmussen et al., 2018). Changes in soil pH have also been shown to correlate with decomposition rates (Leifeld et al., 2013), soil microbial community composition (Lauber et al., 2009), and mineral-organic reactivity (Kleber et al., 2015), among other factors. Thus, soil pH alterations may be a bellwether for soil C effects.

Contrary to our second hypothesis, we did not find evidence of consistent aboveground-belowground correlations for tissue chemistry traits among our study individuals. Plant economic spectrum theory predicts that leaf, stem, and root traits are correlated (Reich, 2014), a prediction that has received support across a range of ecosystems (Freschet et al., 2010; Pérez-Ramos et al., 2012; Shen et al., 2019). However, evidence to the contrary suggests that the root economic spectrum may be multidimensional, potentially leading to a breakdown of aboveground-belowground trait correlations (Bergmann et al., 2017; Weemstra et al., 2016). Thus, the lack of significant couplings between root and stem traits observed under field settings—which is also supported by results obtained from independent greenhouse observations of the natural variant population (Kalluri et al. unpublished data)—indicates a need to collect both aboveground and belowground phenotypic data in future sampling efforts in this system.

Our pilot study suggests that a long-term, genetically diverse plantation can be used to study plant-soil relationships at the scale of individual trees. Yet, we note that we did not actually detect differences related to soil organic matter; a key potential application of these systems. Thus, while our study serves as a proof-of-concept for studying soil in GWAS plantations, more extensive, replicated or longer-term observations are needed to study the impacts of *Populus* genotypes on soil C storage in these systems—which are densely planted with potentially intermeshed root networks. Given the importance of deeper soil layers for soil C storage, future studies should also sample deeper into the soil profile.

## Conclusions

Our results show that genetic diversity plantations can be useful for probing and identifying candidate genotypes and genes based on their effects on key soil biogeochemical properties. Such systems can contain hundreds of individual genotypes, potentially enabling GWAS-scale investigations of soil biogeochemical properties. Our study revealed that the extent of correlation among above-belowground properties across the entire study set is low. Though select correlations between plant and soil properties were detected, we generally found that variability in poplar genotype mattered more than phenotype (e.g. high vs low lignin) for soil biogeochemistry. Our assessments provide new insights into significant negative and positive correlations within a sample type (soil, stem, root) and revealed genotype-specific correlations with associated soil properties, suggesting reciprocal plant-soil impacts that plant genotype and soil depth can influence. We conclude that these effects should be studied more comprehensively considering deeper soil profiles, more genotypes, mineral and particulate associated soil organic C fractions, and microbiome diversity, to facilitate modeling both the productivity and uniformity of aboveground biomass in bioenergy systems, as well as the effects of bioenergy crop production on soil health and carbon sequestration.

## Acknowledgements

This work was supported by the United States Department of Energy (DOE) Center for Bioenergy Innovation (CBI) project. CBI is a Bioenergy Research Center supported by the Office of Biological and Environmental Research in the DOE Office of Science. We thank Eric Pierce for early consultation on the sampling campaign, Sara Jawdy at Oak Ridge National Laboratory for sample aliquoting and shipment to soil service center, Kat Haiby at Poplar Innovations for facilitating field site access, and UGA soil service center for analysis of the study samples. This manuscript has been authored by UT-Battelle, LLC under Contract No. DE-AC05-00OR22725 with the U.S. Department of Energy.

## Author Contributions

MEC contributed to analysis and writing of the manuscript. AEH-W contributed to study design, sampling, analysis, and manuscript writing. KRC contributed to ANOVA analysis of the data. UCK conceived the study and contributed to study design, sampling, analysis, and manuscript writing.

**Suppl. Fig. 1.**
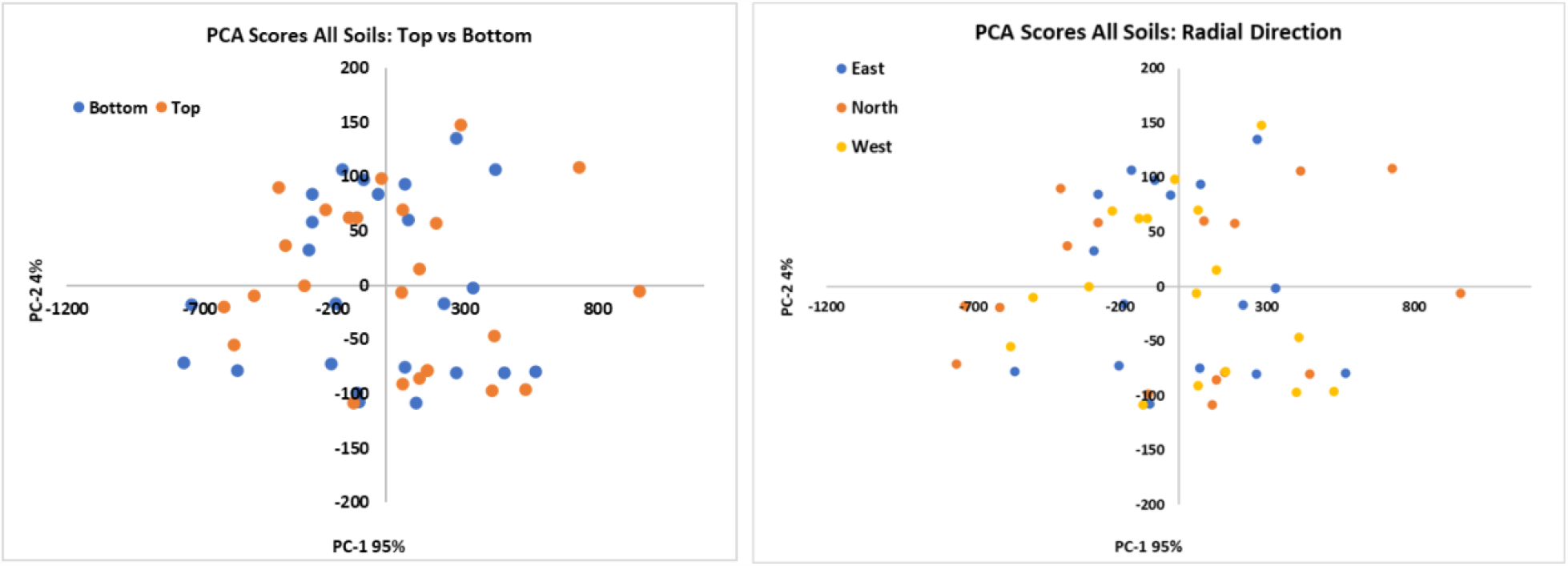
PCA analysis of all soil samples collected from variable depths and radial direction of sampling using compositional analysis traits. PC-1 explains 95% of variance is driven primarily by Ca content while PC-2 corresponds to variation in K content.

**Suppl. Fig. 2.**
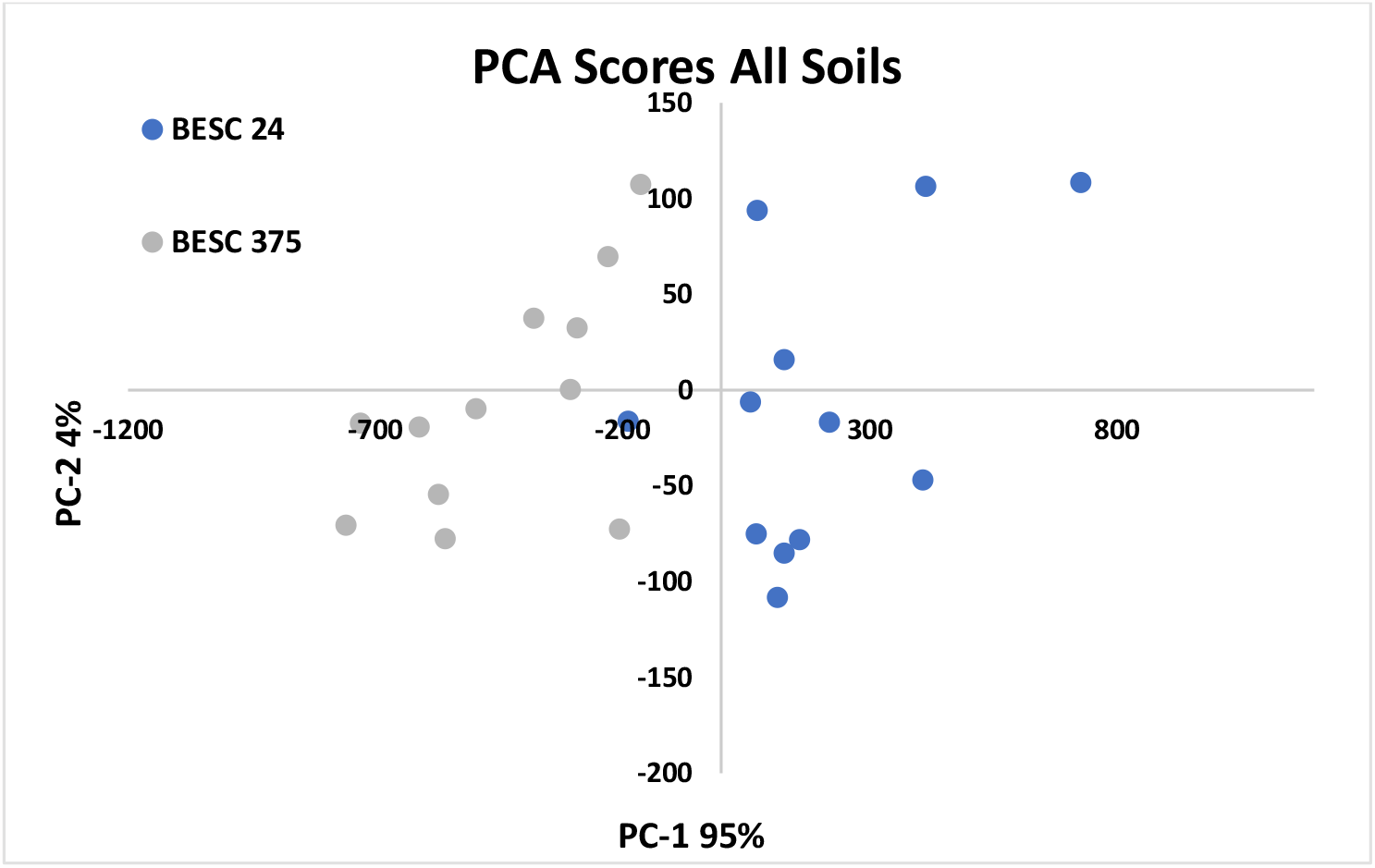
PCA analysis of properties of all soil samples collected from 0-10 cm depth for BESC 24 (low lignin) and BESC 375 (high lignin)

**Suppl. Fig. 3.**
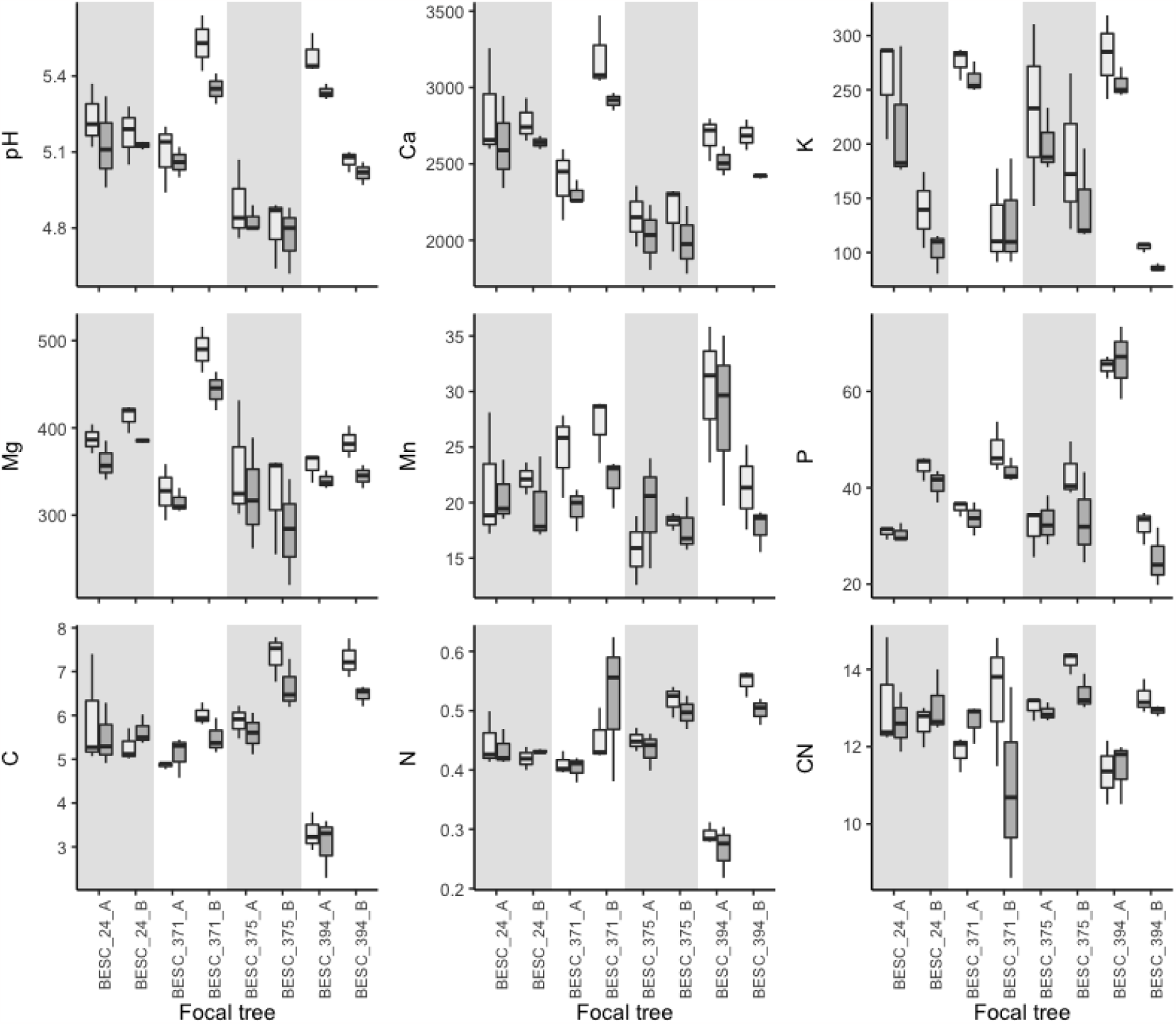
Boxplots showing pH, nutrient concentrations, and C:N in soils sampled at each of eight focal trees belonging to four different *Populus trichocarpa* genotypes (delineated with shading). The two replicates of each genotype (BESC 24, BESC 371, BESC 375, BESC 394) are indicated with the suffix A and B along the x-axis.

**Suppl. Table 1**. Comprehensive correlation matrix presented as a heatmap generated using PCC of all parameters measured for stem, root and soil samples obtained from the four genotypes. See supplemental file.

**Suppl. Table 2.** Variance in soil biogeochemical properties explained (Partial R^2^) by linear models containing either individual trees (n = 3 soil samples per tree) or genotypes (2 trees per genotype, n = 6). Individual trees were significantly related to soil properties (*p* < 0.001) for all measured variables. No statement about statistical significance is made about genotype effects due to insufficient replication at this level of analysis. A significant effect of sample depth is denoted with an asterisk (*) for response variable.

| Variable | Partial $R^2$ | |
| --- | --- | --- |
|  | Individual | Genotype |
| pH* | 0.85 | 0.54 |
| Ca* | 0.79 | 0.53 |
| K* | 0.76 | 0.01 |
| Mg* | 0.72 | 0.27 |
| Mn | 0.51 | 0.26 |
| P* | 0.87 | 0.15 |
| C | 0.86 | 0.19 |
| N | 0.76 | 0.01 |
| C:N | 0.35 | 0.18 |

## References

Bergmann, J., Ryo, M., Prati, D., Hempel, S., & Rillig, M. C. (2017). Root traits are more than analogues of leaf traits: The case for diaspore mass. New Phytologist, 216(4), 1130–1139. https://doi.org/10.1111/nph.14748

Bremner J. M. 1996. Nitrogen-Total, Dumas Method, Chapter 37 in Methods of Soil Analysis. Part 3. Chemical Methods. Soil Science Society of America Book Series No. 5.

Chhetri, H. B., Furches, A., Macaya-Sanz, D., Walker, A. R., Kainer, D., Jones, P., Harman-Ware, A. E., Tschaplinski, T. J., Jacobson, D., Tuskan, G. A., & DiFazio, S. P. (2020). Genome-Wide Association Study of Wood Anatomical and Morphological Traits in Populus trichocarpa. Frontiers in Plant Science, 11, 545748. https://doi.org/10.3389/fpls.2020.545748

Davidson, E. A., & Ackerman, I. L. (1993). Changes in soil carbon inventories following cultivation of previously untilled soils. Biogeochemistry, 20(3), 161–193. https://doi.org/10.1007/BF00000786

Ehrenfeld, J. G., Kourtev, P., & Huang, W. (2001). Changes in Soil Functions Following Invasions of Exotic Understory Plants in Deciduous Forests. Ecological Applications, 11(5), 1287–1300. https://doi.org/10.1890/1051-0761(2001)011[1287:CISFFI]2.0.CO;2

Evans, L. M., Slavov, G. T., Rodgers-Melnick, E., Martin, J., Ranjan, P., Muchero, W., Brunner, A. M., Schackwitz, W., Gunter, L., Chen, J.-G., Tuskan, G. A., & DiFazio, S. P. (2014). Population genomics of Populus trichocarpa identifies signatures of selection and adaptive trait associations. Nature Genetics, 46(10), Article 10. https://doi.org/10.1038/ng.3075

Field, J. L., Richard, T. L., Smithwick, E. A. H., Cai, H., Laser, M. S., LeBauer, D. S., Long, S. P., Paustian, K., Qin, Z., Sheehan, J. J., Smith, P., Wang, M. Q., & Lynd, L. R. (2020). Robust paths to net greenhouse gas mitigation and negative emissions via advanced biofuels. Proceedings of the National Academy of Sciences, 117(36), 21968–21977. https://doi.org/10.1073/pnas.1920877117

Finzi, A. C., Canham, C. D., & Van Breemen, N. (1998). CANOPY TREE–SOIL INTERACTIONS WITHIN TEMPERATE FORESTS: SPECIES EFFECTS ON pH AND CATIONS. Ecological Applications, 8(2), 447–454. https://doi.org/10.1890/1051-0761(1998)008[0447:CTSIWT]2.0.CO;2

Fraterrigo, J. M., Turner, M. G., Pearson, S. M., & Dixon, P. (2005). Effects of Past Land Use on Spatial Heterogeneity of Soil Nutrients in Southern Appalachian Forests. Ecological Monographs, 75(2), 215–230. https://doi.org/10.1890/03-0475

Freschet, G. T., Cornelissen, J. H. C., Van Logtestijn, R. S. P., & Aerts, R. (2010). Evidence of the ‘plant economics spectrum’ in a subarctic flora. Journal of Ecology, 98(2), 362–373. https://doi.org/10.1111/j.1365-2745.2009.01615.x

Garten Jr., C. T., & Wullschleger, S. D. (1999). Soil Carbon Inventories under a Bioenergy Crop (Switchgrass): Measurement Limitations. Journal of Environmental Quality, 28(4), 1359–1365. https://doi.org/10.2134/jeq1999.00472425002800040041x

Gee, G.W. and Bauder, J.W. 1996. Particle Size Analysis: 15-5 Hydrometer Method. Chapter 15 in Methods of Soil Analysis. Part 1-Physical and Mineralogical Methods. SSSA/ASA Book Series No. 5.

Happs, R. M., Bartling, A. W., Doeppke, C., Harman-Ware, A. E., Clark, R., Webb, E. G., Biddy, M. J., Chen, J.-G., Tuskan, G. A., Davis, M. F., Muchero, W., & Davison, B. H. (2021). Economic impact of yield and composition variation in bioenergy crops: Populus trichocarpa. Biofuels, Bioproducts and Biorefining, 15(1), 176–188. https://doi.org/10.1002/bbb.2148

Harman-Ware, A. E., Macaya-Sanz, D., Abeyratne, C. R., Doeppke, C., Haiby, K., Tuskan, G. A., Stanton, B., DiFazio, S. P., & Davis, M. F. (2021). Accurate determination of genotypic variance of cell wall characteristics of a Populus trichocarpa pedigree using high-throughput pyrolysis-molecular beam mass spectrometry. Biotechnology for Biofuels, 14(1), 59. https://doi.org/10.1186/s13068-021-01908-y

Harman-Ware AE, Happs RM, Macaya-Sanz D, Doeppke C, Muchero W, DiFazio SP. 2022. Abundance of Major Cell Wall Components in Natural Variants and Pedigrees of Populus trichocarpa. Front Plant Sci 13: 757810.

Keller, A. B., Brzostek, E. R., Craig, M. E., Fisher, J. B., & Phillips, R. P. (2021). Root-derived inputs are major contributors to soil carbon in temperate forests, but vary by mycorrhizal type. Ecology Letters, 24(4), 626–635. https://doi.org/10.1111/ele.13651

Kissel, D. E., Sonon, L. S., and Cabrera, M. L. (2012). Rapid measurement of soil pH buffering capacity. Soil Sci. Soc. Am. J. 76, 694–699.

Kleber, M., Eusterhues, K., Mikutta, C., Mikutta, R., & Nico, P. S. (2015). Chapter One - Mineral–Organic Associations: Formation, Properties, and Relevance in Soil Environments. In D. L. Sparks (Ed.), Advances in Agronomy (Vol. 130, pp. 1–140). Academic Press. https://doi.org/10.1016/bs.agron.2014.10.005

Kutner, M. H., Nachtsheim, C. J., Neter, J., & Li, W. (2005). Applied Linear Statistical Models (5th Edition). McGraw-Hill.

Lal, R. (2004). Soil Carbon Sequestration Impacts on Global Climate Change and Food Security. Science, 304(5677), 1623–1627. https://doi.org/10.1126/science.1097396

Lauber, C. L., Hamady, M., Knight, R., & Fierer, N. (2009). Pyrosequencing-Based Assessment of Soil pH as a Predictor of Soil Bacterial Community Structure at the Continental Scale. Applied and Environmental Microbiology, 75(15), 5111–5120. https://doi.org/10.1128/AEM.00335-09

Leifeld, J., Bassin, S., Conen, F., Hajdas, I., Egli, M., & Fuhrer, J. (2013). Control of soil pH on turnover of belowground organic matter in subalpine grassland. Biogeochemistry, 112(1), 59–69. https://doi.org/10.1007/s10533-011-9689-5

Lin, G., Craig, M. E., Jo, I., Wang, X., Zeng, D.-H., & Phillips, R. P. (2022). Mycorrhizal associations of tree species influence soil nitrogen dynamics via effects on soil acid–base chemistry. Global Ecology and Biogeography, 31(1), 168–182. https://doi.org/10.1111/geb.13418

Mehlich, A., Determination of P, Ca, Mg, K, Na, and NH4, North Carolina Soil Test Division, Raleigh NC. Mimeo, 1953, p. 16.

Minasny, B., Malone, B. P., McBratney, A. B., Angers, D. A., Arrouays, D., Chambers, A., Chaplot, V., Chen, Z.-S., Cheng, K., Das, B. S., Field, D. J., Gimona, A., Hedley, C. B., Hong, S. Y., Mandal, B., Marchant, B. P., Martin, M., McConkey, B. G., Mulder, V. L., … Winowiecki, L. (2017). Soil carbon 4 per mille. Geoderma, 292, 59–86. https://doi.org/10.1016/j.geoderma.2017.01.002

Muchero, W., Sondreli, K. L., Chen, J.-G., Urbanowicz, B. R., Zhang, J., Singan, V., Yang, Y., Brueggeman, R. S., Franco-Coronado, J., Abraham, N., Yang, J.-Y., Moremen, K. W., Weisberg, A. J., Chang, J. H., Lindquist, E., Barry, K., Ranjan, P., Jawdy, S., Schmutz, J., … LeBoldus, J. M. (2018). Association mapping, transcriptomics, and transient expression identify candidate genes mediating plant–pathogen interactions in a tree. Proceedings of the National Academy of Sciences, 115(45), 11573–11578. https://doi.org/10.1073/pnas.1804428115

Nelson D. W and Sommers, L. E. 1996. Total Carbon by Dry Combustion. Chapter 34 in Method of Soil Analysis. Part 3. Chemical Methods. Soil Science Society of America Book Series No. 5

Paustian, K., Larson, E., Kent, J., Marx, E., & Swan, A. (2019). Soil C Sequestration as a Biological Negative Emission Strategy. Frontiers in Climate, 1. https://doi.org/10.3389/fclim.2019.00008

Paustian, K., Lehmann, J., Ogle, S., Reay, D., Robertson, G. P., & Smith, P. (2016). Climate-smart soils. Nature, 532(7597), Article 7597. https://doi.org/10.1038/nature17174

Pérez-Ramos, I. M., Roumet, C., Cruz, P., Blanchard, A., Autran, P., & Garnier, E. (2012). Evidence for a ‘plant community economics spectrum’ driven by nutrient and water limitations in a Mediterranean rangeland of southern France. Journal of Ecology, 100(6), 1315–1327. https://doi.org/10.1111/1365-2745.12000

Rasmussen, C., Heckman, K., Wieder, W. R., Keiluweit, M., Lawrence, C. R., Berhe, A. A., Blankinship, J. C., Crow, S. E., Druhan, J. L., Hicks Pries, C. E., Marin-Spiotta, E., Plante, A. F., Schädel, C., Schimel, J. P., Sierra, C. A., Thompson, A., & Wagai, R. (2018). Beyond clay: Towards an improved set of variables for predicting soil organic matter content. Biogeochemistry, 137(3), 297–306. https://doi.org/10.1007/s10533-018-0424-3

Rasse, D. P., Rumpel, C., & Dignac, M.-F. (2005). Is soil carbon mostly root carbon? Mechanisms for a specific stabilisation. Plant and Soil, 269(1), 341–356. https://doi.org/10.1007/s11104-004-0907-y

Reich, P. B. (2014). The world-wide ‘fast–slow’ plant economics spectrum: A traits manifesto. Journal of Ecology, 102(2), 275–301. https://doi.org/10.1111/1365-2745.12211

Sanderman, J., Hengl, T., & Fiske, G. J. (2017). Soil carbon debt of 12,000 years of human land use. Proceedings of the National Academy of Sciences, 114(36), 9575–9580. https://doi.org/10.1073/pnas.1706103114

Shen, Y., Gilbert, G. S., Li, W., Fang, M., Lu, H., & Yu, S. (2019). Linking Aboveground Traits to Root Traits and Local Environment: Implications of the Plant Economics Spectrum. Frontiers in Plant Science, 10, 1412. https://doi.org/10.3389/fpls.2019.01412

Sokol, N. W., & Bradford, M. A. (2019). Microbial formation of stable soil carbon is more efficient from belowground than aboveground input. Nature Geoscience, 12(1), 46–53. https://doi.org/10.1038/s41561-018-0258-6

Sun, T., Hobbie, S. E., Berg, B., Zhang, H., Wang, Q., Wang, Z., & Hättenschwiler, S. (2018). Contrasting dynamics and trait controls in first-order root compared with leaf litter decomposition. Proceedings of the National Academy of Sciences, 115(41), 10392–10397. https://doi.org/10.1073/pnas.1716595115

Vesterdal, L., Schmidt, I. K., Callesen, I., Nilsson, L. O., & Gundersen, P. (2008). Carbon and nitrogen in forest floor and mineral soil under six common European tree species. Forest Ecology and Management, 255(1), 35–48. https://doi.org/10.1016/j.foreco.2007.08.015

Villarino, S. H., Pinto, P., Jackson, R. B., & Piñeiro, G. (2021). Plant rhizodeposition: A key factor for soil organic matter formation in stable fractions. Science Advances, 7(16), eabd3176. https://doi.org/10.1126/sciadv.abd3176

von Lützow, M., Kögel-Knabner, I., Ekschmitt, K., Flessa, H., Guggenberger, G., Matzner, E., & Marschner, B. (2007). SOM fractionation methods: Relevance to functional pools and to stabilization mechanisms. Soil Biology and Biochemistry, 39(9), 2183–2207. https://doi.org/10.1016/j.soilbio.2007.03.007

Weemstra, M., Mommer, L., Visser, E. J. W., van Ruijven, J., Kuyper, T. W., Mohren, G. M. J., & Sterck, F. J. (2016). Towards a multidimensional root trait framework: A tree root review. New Phytologist, 211(4), 1159–1169. https://doi.org/10.1111/nph.14003

Yang, X., Liu, D., Lu, H., Weston, D. J., Chen, J.-G., Muchero, W., Martin, S., Liu, Y., Hassan, M. M., Yuan, G., Kalluri, U. C., Tschaplinski, T. J., Mitchell, J. C., Wullschleger, S. D., & Tuskan, G. A. (2021). Biological Parts for Plant Biodesign to Enhance Land-Based Carbon Dioxide Removal. BioDesign Research, 2021. https://doi.org/10.34133/2021/9798714

